# Automated High-Throughput Light-Sheet Fluorescence Microscopy of Larval Zebrafish

**DOI:** 10.1101/330639

**Authors:** Savannah L. Logan, Christopher Dudley, Ryan P. Baker, Michael J. Taormina, Edouard A. Hay, Raghuveer Parthasarathy

**Author notes:** Equal contributors (Savannah L. Logan, Christopher Dudley, Ryan P. Baker).

## Abstract

Light sheet fluorescence microscopy enables fast, minimally phototoxic, three-dimensional imaging of live specimens, but is currently limited by low throughput and tedious sample preparation. Here, we describe an automated high-throughput light sheet fluorescence microscope in which specimens are positioned by and imaged within a fluidic system integrated with the sheet excitation and detection optics. We demonstrate the ability of the instrument to rapidly examine live specimens with minimal manual intervention by imaging fluorescent neutrophils over a nearly 0.3 mm^3^ volume in dozens of larval zebrafish. In addition to revealing considerable inter-individual variability in neutrophil number, known previously from labor-intensive methods, three-dimensional imaging allows assessment of the correlation between the bulk measure of total cellular fluorescence and the spatially resolved measure of actual neutrophil number per animal. We suggest that our simple experimental design should considerably expand the scope and impact of light sheet imaging in the life sciences.

## INTRODUCTION

Light sheet fluorescence microscopy (LSFM) is a powerful tool for examining the three-dimensional structural and temporal dynamics of living systems. In LSFM, a thin sheet of laser light excites fluorophores in a sample. Scanning the sample through the sheet enables fast, three-dimensional imaging with low phototoxicity, high resolution, and a wide field of view (1–5). Imaging with LSFM has enabled numerous studies of embryonic development (6,7), neural activity (8), microbial dynamics (9–11), and other phenomena. A large body of work has focused on improving the optical capabilities of light sheet imaging, for example using structured illumination (12,13), multiple lens pairs (6), two-photon excitation (14), and other techniques. However, current light sheet fluorescence microscopes have significant constraints related to throughput and sample handling that, we argue, have placed much greater limitations on their scientific utility than issues of spatial or temporal resolution. The majority of existing light sheet fluorescence microscopes, both commercial and non-commercial, are designed to hold a single specimen. A few instruments (including one from the authors of this paper) can hold up to six specimens for sequential imaging. Moreover, light sheet fluorescence microscopy typically requires extensive sample preparation and manual sample mounting, most commonly by embedding specimens in agarose gels. Examination of large numbers of specimens is therefore slow and difficult, which is especially important given the high level of inter-individual variability found in many complex biological processes. Increasing the pace of insights into developmental biology, multicellular biophysics, or microbial community structure will require faster and simpler acquisition of three-dimensional imaging datasets. To date, there exists only one report of an automated light sheet microscope that makes use of fluidic positioning of live animals (15); its throughput (specimens per hour) is not stated, and though its ability to image larval zebrafish is clear, the total number of animals examined was only twelve. In contrast, automated, high-throughput methods have been integrated with other types of microscopes, including confocal microscopes (16–18), about which we comment further in the Discussion. Our system adopts and builds upon these, especially the confocal-based setup of Ref. (16).

To address the issues described above, we developed a light sheet fluorescence microscope capable of automated, high-throughput imaging of live specimens. Our instrument uses fluidic control and image-based registration to rapidly but precisely position specimens for light sheet scans and subsequently remove them from the imaging area. We characterize the optical quality of our instrument, and demonstrate its capabilities by rapidly imaging immune cells in dozens of larval zebrafish. While the spatial resolution of our microscope does not equal that of current “low-throughput” light sheet microscopes, it is more than sufficient for determining cellular distributions. Moreover, we argue that the tradeoff of lower resolution for higher throughput is worthwhile given the large variance in most biological datasets.

We illustrate the utility of the instrument by imaging neutrophils in dozens of larval zebrafish. Neutrophils are an important and highly dynamic cell type of the innate immune system. These cells migrate to sites of damage or infection and recognize and kill pathogenic microbes (19). Zebrafish are a well recognized model organism for studying neutrophil responses (20,21), and prior work has uncovered changes in neutrophil distributions in response to wound-induced chemotaxis (22), drugs that lead to symptoms similar to human enterocolitis (23), and stimulation by gut microbes or bacterial lipopolysaccharide (LPS) (24,25), to list a few of many examples. Quantifing even basic properties such as neutrophil abundances over large extents for the dozens of specimens required given the high variance between individuals often necessitates slow confocal imaging or painstaking histological sectioning or gut dissection (24–26). These sorts of studies will be greatly facilitated by instruments that can provide large-scale 3D imaging of automatically detected and imaged specimens.

We show, as expected, a high degree of variation between fish in total neutrophil number and an increase in its mean value following LPS exposure, as well as spatial clustering of neutrophils in two distinct regions near the swim bladder. While our instrument is optimized for imaging of larval zebrafish, the design could easily be modified for rapid imaging of a wide range of biological and non-biological samples, which should broaden the impact of light sheet microscopy in a variety of fields.

## RESULT

### Instrument design

The light sheet portion of the microscope closely follows the design of Keller et al (1); a rapidly-scanned galvanometer creates a sheet of light for fluorescence excitation, and emitted light is captured by a camera perpendicular to the plane of the sheet (Fig. 1A). In conventional light sheet fluorescence microscopes, gel-mounted specimens are introduced vertically in between horizontal lenses. To achieve high throughput, we use a continuous fluidic path through plastic tubing and glass capillaries for transport as well as imaging, detailed below. If the fluidic path were oriented vertically, specimens would gravitationally drift during imaging. Therefore, we adopted a geometry in which specimens are transported horizontally and the sheet plane is vertical (Fig. 1A,B). To allow this arrangement, we designed an elongated sample chamber with windows oriented below and perpendicular to the sample (Fig. 1B,C). Before entering the imaging chamber, specimens flow through a system of 0.7 mm diameter plastic tubing at a typical flow rate of 1 ml/min, or 4 cm/sec (Fig. 1). Flow speed and direction are controlled by a syringe pump (Fig. 1); see Supplemental Methods for a parts list and descriptions.

**Figure 1.**
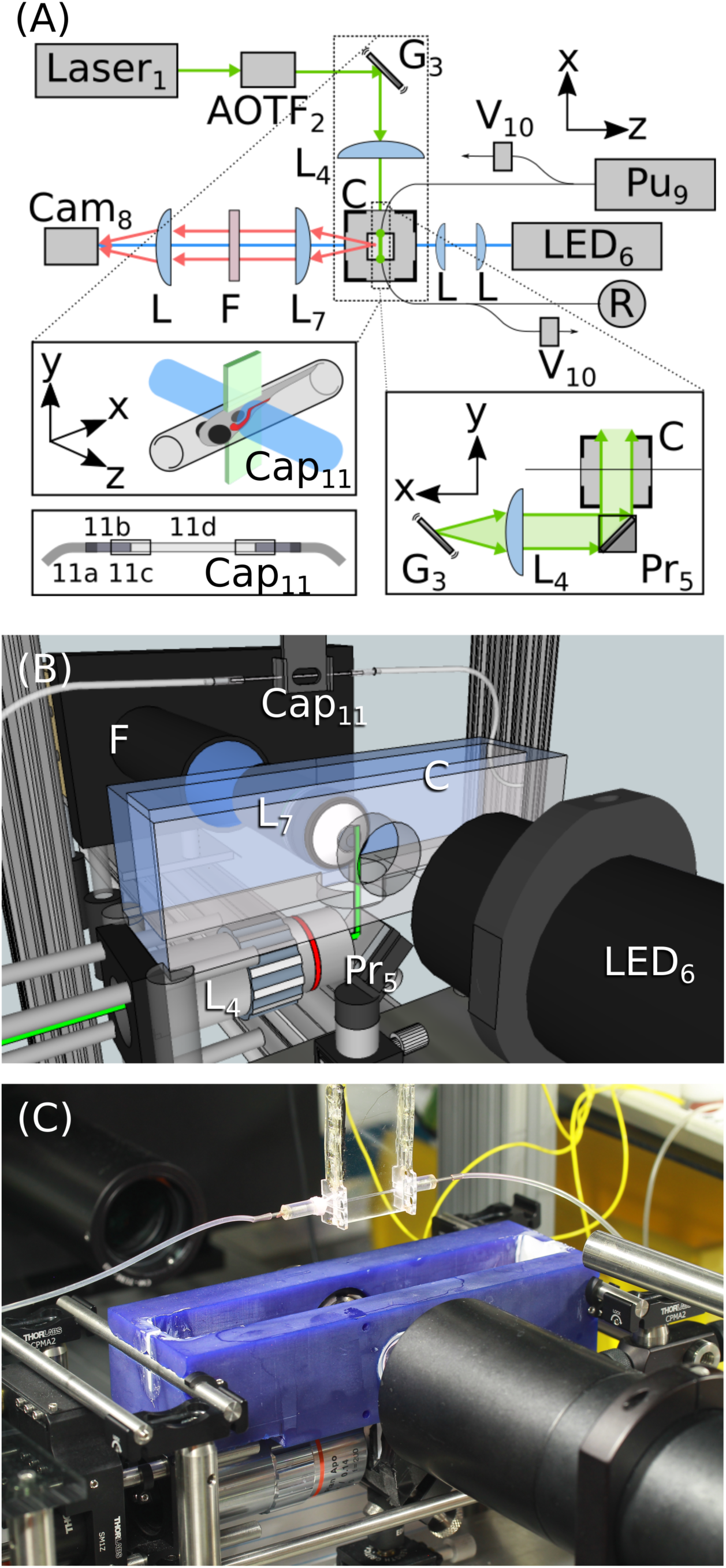
Instrument design. **(A)** Schematic of the instrument design, with labels corresponding to the parts list in Table 1. See also Supplementary Movie 1. The excitation laser line is selected by an acousto-optic tunable filter (AOTF_2_), then directed to a galvanometer mirror (G_3_) and objective lens (L_4_) to create a time-averaged sheet of light in the sample chamber (C) via a prism (Pr_5_). Specimens flow through a system of tubing controlled by a syringe pump (Pu_9_) and valves (V_10_) and are automatically positioned in a square-walled capillary (Cap_11_) for imaging. Bright field images are used for positioning the sample and are illuminated with an LED (LED_6_). After imaging, specimens are directed into a reservoir (R). **(B)** Schematic of the imaging area. The 3D-printed sample chamber (C), prism (Pr_5_), and imaging capillary (Cap_11_) are apparent. **(C)** Photograph of the imaging area corresponding to the schematic in (B).

Inside the imaging chamber, specimens flow into a square-walled glass capillary in front of the imaging objective where they are automatically detected by bright field microscopy. Specimens are rapidly stopped using computer-controlled valves on either side of the imaging chamber, with a precision of approximately 1 mm in position, comparable to the length of a larval zebrafish. Fine positioning is performed by iterated movement of the capillary by a computer-controlled stage, brightfield imaging, and comparison of images with a previously assembled image library (see Methods) (Fig. 2 A,B). The travel range of the capillary on the stage allows movement of up to 30 mm in the x-direction. Like many studies, ours make use of larval zebrafish as a model organism; strong features such as eyes and the swim bladder enable straightforward correlation-based registration, with a precision of about 20 µm as described below. This approach should be applicable to any specimen with a roughly sterotypical anatomy. The specimen is not rotated about the tube axis; we comment further on this in the Discussion section.

Once positioned, specimens are automatically imaged using LSFM. The imaging chamber has sufficient depth, 35 mm, that the capillary can be scanned through the sheet by a motorized stage. In our setup, repeated scans with up to three excitation wavelengths are possible; this is limited simply by the number of available laser lines. The precision of the automated positioning enables scans to be taken of particular regions, for example the larval gut, as shown below. After imaging, specimens flow into a collection reservoir and subsequent specimens are automatically positioned for imaging. We provide a movie of the instrument in operation as Supplementary Video S1. A complete parts list is provided as Table 1, in Supplemental Methods.

### Optical quality

Our instrument uses glass capillaries for specimen mounting, rather than more conventional gel embedding. The square cross-section of these capillaries should lead to less distortion than more common cylindrical capillaries. To assess the optical quality of our setup, we measured the point spread function (PSF) by imaging 28 nm diameter fluorescent microspheres dispersed in oil in these capillaries (see Methods for details). The diffraction-limited width of the particles in the sheet plane (*xy*), assessed as the standard deviation of a Gaussian function fit to the particle’s intensity profile, is 0.6 µm, and the width along the detection axis (*z*) is 3.4 ± 0.6 µm, consistent with the expected sheet thickness of our setup (Fig. 2 C,D).

**Figure 2.**
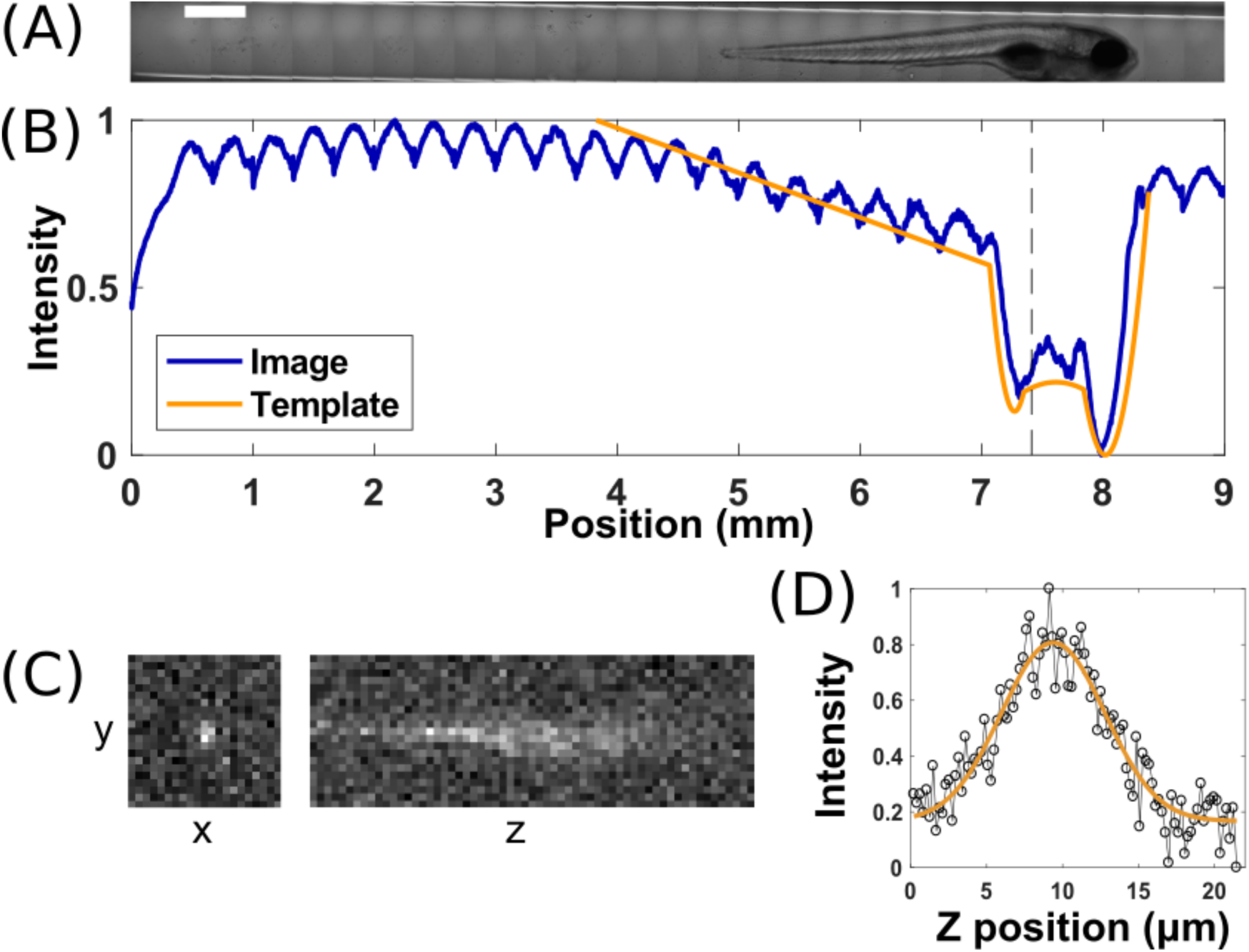
Specimen positioning and image quality. (A) Composite brightfield image of a larval zebrafish positioned in a glass capillary. Scale bar: 50 µm. (B) Normalized intensity averaged along the short axis of the brightfield image, and the intensity of the template image that best matches the fish in (A). Cross-correlation with the template is used to automatically position the fish for light sheet fluorescence imaging. (C) Light sheet fluorescence images of a 28 nm diameter fluorescent microsphere, showing *x*-*y* and *z*-*y* planes centered on the particle. (D) Line-scan of intensity along the detection axis (*z*) through a fluorescent microsphere, with a Gaussian fit showing a width of approximately 3 µm.

### Data collection capabilities

Using this system, we can image approximately 30 larval zebrafish per hour, obtaining from each a 666 x 431 x 1060 µm (*x, y, z*) three-dimensional scan, a marked improvement over manual mounting and imaging that, even by a skilled researcher, is limited to about 5 fish per hour. The triggering accuracy is about 90%, with roughly 10% of detected objects being bubbles or debris that are easily identified after imaging. On average, 81% of larval fish are automatically positioned correctly in front of the imaging objective. The remaining 19% correspond to multiple fish being in the field of view, or other positioning errors. Importantly, the majority of the run time of the instrument is spent obtaining light sheet fluorescence images, and is not dominated by specimen positioning. In the batch of *N*=41 fish whose neutrophil distributions were imaged in experiments described below, for example, the flow, detection, and positioning of the specimens occupied only approximately 30 seconds per fish. In the limit of zero imaging time (e.g. for very bright signals or small regions of interest), the system could therefore record data from up to about 120 specimens per hour in the absence of triggering or positioning errors, or about 90 specimens per hour with the present system performance.

### Neutrophils in larval zebrafish

To demonstrate the capabilities of the instrument, we imaged fluorescent neutrophils in larval zebrafish at 5 days post-fertilization (dpf), focusing especially on the number of these immune cells and their distribution near the anterior of the intestine. Specifically, these were fish engineered to express green fluorescent protein (GFP) under the promoter myeloperoxidase, an enzyme primarily produced in neutrophils (27).

### Positioning accuracy

To assess the positioning accuracy of our instrument, we performed automated scans of larval zebrafish with GFP-expressing neutrophils, and then reloaded the same fish and rescanned them. Changes in the neutrophil positions are due to both imaging error and to motion of the neutrophils during the intermediate time. (Neutrophils are highly motile cells, crawling through tissue and also entering or leaving tissue via the bloodstream.) For five twice-scanned fish, we manually identified three neutrophils that were unambiguously the same in each scan (i.e. not newly entered or departed). For these neutrophils, the within-fish standard deviations of the changes in position provide a measure of the biological variation, e.g. from neutrophil motion. These were 5.3 ± 3.5 µm, 11.6 ± 9.0 µm, and 40.0 ± 36.4 µm, for *x, y*, and *z*, respectively, where *x* is the flow direction and *z* is perpendicular to the light sheet. The standard deviation between fish of the mean neutrophil positions provides a measure of the instrumental variation, e.g. from imperfect positioning. These were 18.9 ± 6.7 µm, 56.1 ± 19.9 µm, and 52.6 ± 18.6 µm for *x, y*, and *z*, respectively. Along the capillary axis, therefore, the fish positioning is reproducible to within about 20 µm. The larger variance in *y* and *z* is expected, as we are not explicitly detecting location along these axes, and because there can be specimen rotation about the *x*-axis. Overall, therefore, global positioning uncertainty is on the order of a cell diameter for cells such as neutrophils.

### Neutrophil number and variance

In total, we imaged 41 fish, obtaining from each a single 666 x 431 x 1060 µm three-dimensional image in which neutrophils were readily evident (Fig. 3A and Supplementary Movie S2). Brightfield images are captured and saved prior to fluorescence imaging; a set of four such images, demonstrating their appearance and variance, are provided as Supplemental Figure 1. The strong GFP signal enabled automated identification of neutrophils by standard segmentation methods. Corroborating previous work done by manual dissection of zebrafish (24), we found a high degree of variation in neutrophil number between specimens, with the standard deviation being 30% of the mean (Fig. 3B). Furthermore, we found that neutrophils tend to cluster in two distinct regions: adjacent to the swim bladder on both the anterior and posterior (Fig. 3C). Notably, the total GFP intensity in a fish is weakly correlated with the number of neutrophils, with a coefficient of determination R^2^ = 0.4, indicating that a simple measure of overall brightness, as could be assessed without three-dimensional microscopy, would provide a poor diagnostic of the actual abundance of immune cells (Fig. 3D). We also compared neutrophil counts determined from two-dimensional maximum intensity projections of the full three-dimensional image stacks, mimicking images that would be obtained from simple widefield fluorescence microscopy. This yielded a number of neutrophils that was on average 0.76 ± 0.02 of that from the three-dimensional counts, indicating as expected that some neutrophils are behind others in the three-dimensional space of the specimen, and hence require three-dimensional imaging for accurate assessment (Supplemental Figure 2).

**Figure 3.**
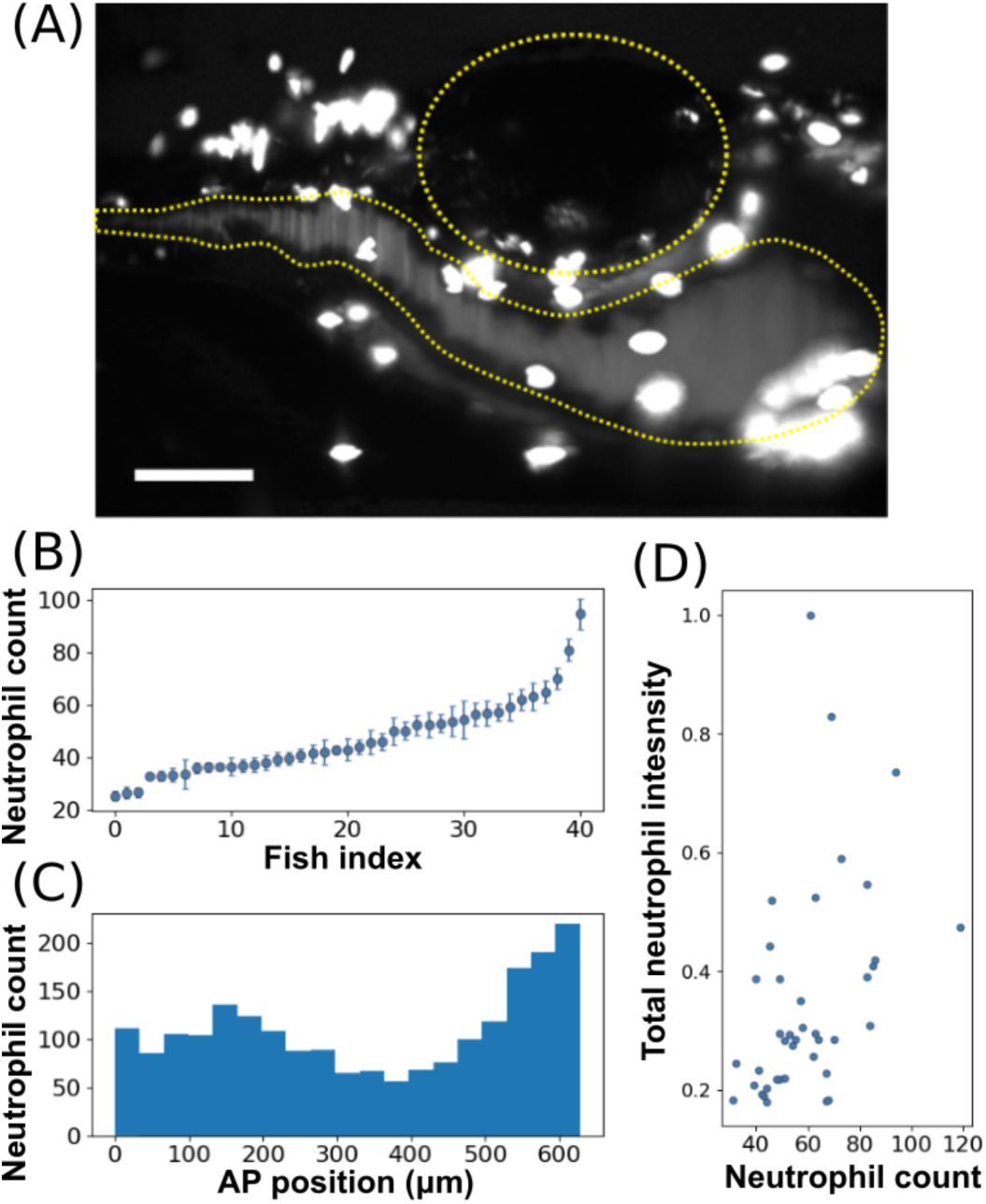
Imaging neutrophils in larval zebrafish. (A) A maximum intensity projection of a three-dimensional light sheet fluorescence image of GFP-expressing neutrophils near the intestine of a 5 dpf larval zebrafish. The 3D scan is provided as Supplemental Movie 2. The intestine and swim bladder are roughly outlined by the yellow dotted lines. Scale bar: 100 µm. (B) The total number of neutrophils in each fish; the x-axis is ordered by neutrophil count. (C) Neutrophil count along the anterior-posterior dimension, summed over all fish examined (*N*=41). The x-axis corresponds approximately to the horizontal range of (A). (D) The total intensity of the detected neutrohils per fish vs the total number of neutrophils in that fish. The two measures are weakly correlated with a coefficient of determination R^2^ = 0.4.

In addition, we examined neutrophil number in response to exposure of larval zebrafish to soluble LPS, the major component of the outer membrane of gram negative bacteria, known from earlier studies to stimulate an inflammatory immune reaction (25,28). As above, we detected and scanned transgenic zebrafish with GFP-expressing neutrophils, exposed to LPS in their flask water at a concentration of 150 µg/ml for 0 (control), 2, and 24 hours prior to imaging. As above, neutrophil number shows a large degree of variability. At 5 dpf, we detected 85.0 ± 17.0 (*N* = 13) and 87.6 ± 27.6 (*N* = 14) (mean ± standard deviation) neutrophils in fish subjected to no LPS and LPS for 2 hours, respectively; indicating no discernible change. At 6 dpf, we detected 95.1 ± 23.4 (*N* = 21) and 114.2 ± 38.1 (*N* = 19) in the control and 24-hour LPS-treated fish, respectively, indicating an increase in the mean neutrophil count by a factor of 1.2 ± 0.1.

**Figure 4.**
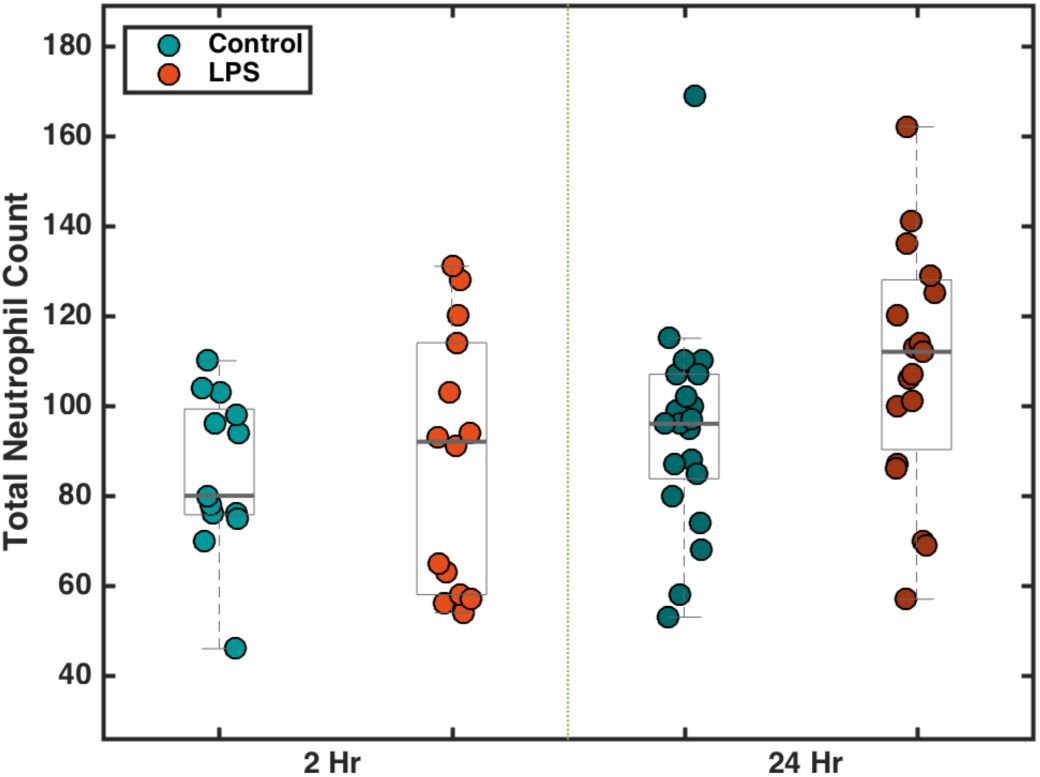
Neutrophil counts after exposure to LPS. The number of neutrophils counted from light sheet fluorescence images of larval zebrafish after exposure to 150 µg/ml LPS for 0 (control), 2, or 24 hours. At 2 hours post-exposure, there was no discernible difference between the LPS-treated group and the control group. At 24 hours post-exposure, the LPS group displayed an increase in mean neutrophil count of 1.2 ± 0.1.

## DISCUSSION

In recent years, many instruments for high content and high throughput imaging of small organisms, such as embryonic and larval zebrafish, have been developed (15–18,29–32), and have demonstrated their utility for fast, accurate imaging and screening. Our setup has similarities and differences to several of these, and advantages and disadvantages for particular applications. Several instruments are designed to image specimens held in multi-well plates (29–32), including commercial instruments such as the ImageXpress High-Content Screening System (33) and the Acquifer Imaging Machine (34). These plate-based systems enable integration with standard specimen containment, and also facilitate automated delivery of chemicals or other perturbations. Imaging is typically provided by widefield microscopy, which is rapid and simple but which prohibits three-dimensional imaging. While confocal microscopy is in principle possible, the thickness of standard plate bottoms would make this difficult except for specialized setups. Three-dimensional confocal fluorescence microscopy is, however, attained in automated instruments such as those of Refs. (16) and the commercial VAST device (35), which use capillary- and tubing-based automated specimen handling and positioning. While successful, and enabling a wide range of studies, there are applications for which the advantages of light sheet microscopy compared to confocal microscopy, namely its rapid speed and low phototoxicity (1–5,36), are important, such as the imaging of active cells in dynamic environments like the intestine. The requirement of orthogonal accessible light paths for light sheet imaging make its integration with existing high-throughput instruments non-trivial, motivating the work presented here. Our design of a relatively freestanding capillary amid a vertical excitation sheet and horizontal detection axis is effective, but it need not be the only solution. For example, one could imagine integration with single-lens-based light sheet techniques (37,38), and it will be interesting to see if such schemes are developed. We note that our device will have lower throughput than plate-based instruments, and will not integrate with commercial confocal microscopes as existing tubing-based methods do, but rather will enhance studies for which light sheet fluorescence microscopy allows insights into dynamic biological phenomena that are otherwise unattainable.

While our microscope is optimized for rapid imaging of larval zebrafish, the design is general and opens numerous possibilities for imaging a wide range of specimens, such as organoids, drosophilia embryos, small marine invertebrates, and more, with the appropriate expansion of the positioning image library.

Our design does not rotate the specimen about the travel axis, a design choice that has advantages and disadvantages for applications. It is certainly possible to vary the specimen orientation by rotating the glass capillary, as, for example, in the VAST instrument (35). Rotation would enhance image quality, both by allowing the selection of particular orientations with minimal sheet distortion and by enabling the fusion of multiple views to gain isotropic resolution. We note, moreover, that variation in image quality due to the uncontrolled orientations of zebrafish likely contributes, along with true biological variability, to the total scatter in neutrophil counts (Figs. 3, 4). The ability to examine a large number of specimens allows averaging over both sources of variation. Though it could be implemented, rotational positioning of each animal takes time and, at least in our design, significantly reduces the throughput of the instrument. Furthermore, we believe that many useful applications, e.g. the neutrophil counting demonstrated here, can be realized without specimen rotation, and that the simplicity of the setup as presented can hopefully foster widespread adoption. We note that rotation can be decoupled from the imaging area, as in Ref. (35), so that one specimen can be oriented while another is imaged, preventing a reduction in throughput. Such additions to the instrument described, though carrying a cost of greater complexity, may be worthwhile.

While the present design provides only a single three-dimensional image of each specimen, we envision future integration of a closed-loop fluidic circuit, with which specimens can be automatically loaded, imaged, and circulated repeatedly, allowing for high-throughput acquisition of multiple snapshots of the same specimen over time. In general, tackling the challenge of automated, high-throughput specimen handling will allow the technique of light sheet fluorescence microscopy to maximize its impact on the life sciences.

## METHODS

### Hardware

The majority of the instrument was constructed with off the shelf parts. Custom parts were either laser cut from acrylic sheets or were 3D printed (see Parts List).

Fluorescence excitation is provided by various solid state lasers, selected by an acousto-optic tunable filter (AOTF, AA Opto-electronic) for coupling into a fiber launch to a galvanometer mirror (Cambridge Technology), which oscillates with a triangular waveform at 1 kHz to sweep the beam into a sheet. An objective lens (Mitutoyo 5X) and a prism route the sheet to the water-filled sample chamber where it intersects the specimen (Fig. 1). Detection is provided by a 20x water immersion objective (Olympus 20XW), mounted in the side of the chamber and sealed with an o-ring, its corresponding tube lens, and an sCMOS camera (Hamamatsu Orca Flash 4.0). Exposure times were 25 ms for all experiments presented here. Instrument control software was written in MATLAB.

### Ethics statement

All zebrafish experiments were carried out in accordance with protocols approved by the University of Oregon Institutional Animal Care and Use Committee (39).

### Zebrafish husbandry

The zebrafish line *Tg[BACmpo:gfp]* (27) was used for neutrophil imaging. Larval zebrafish were raised at a density of one embryo per milliliter and kept at a constant temperature of 28°C. Embryos were not fed during experiments and were sustained by their yolks.

### Sample preparation

Larval zebrafish were placed in dishes containing sterile embryo media with 0.05% methylcellulose and anaesthetized using 240 µg/ml tricaine methanesulfonate. This anaesthetic concentration is higher than the standard dosage, but was necessary likely because of permeation through the plastic tubing. Specimens are initially loaded into the tubing system by manual aspiration using a syringe connected to the opposite end of the tubing, maintaining a spacing between specimens of approximately 6 inches. Fish are pulled into the tubing head-first to ensure high and consistent flow speeds. (Fish travel better forwards than backwards). During experiments, larvae flowed through the tubing and were automatically stopped and positioned by a syringe pump and a series of valves. After imaging, larvae flowed into a dish containing sterile embryo medium.

### LPS Treatment

A filter-sterilized LPS (*E. coli* serotype 0111:B4, Sigma) solution was injected into flasks containing 15 zebrafish larvae at 5 dpf for a final concetration of 150 µg/ml of LPS in each flask. Fish were incubated in the LPS solution for 2-24 hours as indicated in the text, after which they were removed from the solution and imaged.

### Imaging Procedure

For live imaging, each larval zebrafish flows through 0.7 mm inner diameter silicone tubing to a 50 mm long section of round 0.7 mm inner diameter, square exterior cross section, glass tubing in the specimen chamber. The contrast between the specimen and background in brightfield images is used to detect the specimen, stop the pump, and close off tubing using pinch valves to prevent specimen drift. Detecting the specimen and halting the flow places the specimen within three to five millimeters of the desired location. Fine positioning of the specimen is performed by translating the specimen along the capillary axis (“x”) in roughly 30 steps of 0.3 mm size, capturing a brightfield image at each position, tiling the brightfield images into one large image, and comparing the result with a previously assembled “library” image. The comparison can be done in one of two ways, which yield indistinguishable positioning accuracies, both of which begin by averaging the image intensity along the “y” direction, giving a one-dimensional intensity profile. The profile is then compared with the profile derived from average library images either (i) by cross-correlation, in which one profile is offset in position and then multiplied by the other profile, and integrated (summed), giving a correlation value as a function of offset value, or (ii) the location of the profile’s intensity minimum is determined, and compared to the location of the average library profile’s minimum. In (i) the peak in the correlation function and in (ii) the difference between the minima gives the displacement between the profile and the library profile, and therefore indicates where the specimen should be precisely positioned. Following positioning, bright field illumination is switched off and the desired laser wavelength is selected by the AOTF. The *xyz* arm is then scanned perpendicular to the sheet plane, generating a 3D image stack. The scan can be repeated for another region or another wavelength before the pump is directed to send the specimen out of the chamber, and bring in the next specimen.

### Point Spread Function

For measurements of the point spread function, 28 nm diameter fluorescent carboxylate-modified polystyrene spheres (Thermo Fisher cat. #F8787), with peak excitation and emission wavelengths 505 and 515 nm, respectively, were dispersed in oil with a similar index of refraction as water (Zeiss Immersol W 2010) inside the imaging capillaries. Three-dimensional scans were taken, with a z-spacing of 0.5 µm.

### Image-based neutrophil quantification

Neutrophils were detected from image stacks using custom code written in Python, involving a coarse and fine thresholding. First, the 3D stack was thresholded using a low intensity value followed by the morphological operations of closing and erosion with a range of 1 pixel (40). Connected above-threshold pixels were identified as objects using scikit-image’s label function. Then, each detected object was further thresholded using an Otsu filter and labeled again. This re-thresholding process was repeated twice, and objects below 3 µm^3^ in volume were discarded as unphysical. The total intensity within each identified object represents the total fluorescence intensity of that neutrophil. For a full 3D dataset, the processing takes 1-2 minutes.

### Raw data

Raw image data, roughly 150 GB in total size, is available from the authors on request.

## ACKNOWLEDGEMENTS

We thank Rose Sockol, Kyleah Murphy, and the University of Oregon Zebrafish Facility staff for fish husbandry, and Karen Guillemin and Brandon Schlomann for useful comments and conversations. Research reported in this publication was supported in part by the National Science Foundation (NSF) under award 1427957 (to RP), by the the M. J. Murdock Charitable Trust, and by the National Institutes of Health (NIH) as follows: by the National Institute of General Medical Sciences under award number P50GM098911 and by the National Institute of Child Health and Human Development under award P01HD22486, which provided support for the University of Oregon Zebrafish Facility. The content is solely the responsibility of the authors and does not represent the official views of the NIH, NSF, or other funding agencies.

## SUPPORTING INFORMATION CAPTIONS

**Supplemental Movie 1.** Video of the automated, high-throughput light sheet microscope in operation. After a larval zebrafish, flowing through tubing, is detected and positioned, 3D scans in two colors in two different regions, are obtained. The final image (0:57-1:03) is of tdTomato-labeled bacteria and GFP-labeled neutrophils.

**Supplemental Movie 2.** A light sheet fluorescence scan through the anterior intestine of a larval zebrafish with GFP-expressing neutrophils evident as very bright objects above autofluorescent background. The total image depth is 475 µm (2.5 µm/slice x 190 slices). The maximum intensity projection of this scan is shown in Figure 2E.

**Supplemental Figure 1.** Sample brightfield images of four larval zebrafish, captured and saved prior to light sheet fluorescence imaging. Notably, the orientation is random in the capillary, with a tendancy toward either “gut down” or “gut up.” Scale bar: 100 µm.

**Supplemental Figure 2.** The number of neutrophils in 67 larval zebrafish, assessed from two-dimensional maximum intensity projections and from the full three-dimensional light sheet fluorescence scans. The former is 0.76 ± 0.02 of the latter, indicating that three-dimensional imaging is necessary to capture all the cells in a three-dimensional volume.

